# The post-translational modification SUMO affects TDP-43 phase separation, compartmentalization, and aggregation in a zebrafish model

**DOI:** 10.1101/2022.08.14.503569

**Authors:** Cindy Maurel, Natalie M. Scherer, Alison Hogan, Andres Vidal-Itriago, Emily K. Don, Rowan Radford, Tyler Chapman, Stephen Cull, Patrick Vourc’h, Roger Chung, Albert Lee, Marco Morsch

## Abstract

TDP-43 is a nuclear RNA-binding protein that can undergo liquid-liquid phase separation (LLPS) and forms pathological insoluble aggregates in frontotemporal dementia and amyotrophic lateral sclerosis (ALS). Perturbations of TDP-43 function are linked to mislocalization and neurodegeneration. By studying TDP-43 *in vivo*, we confirmed for the first time that TDP-43 undergoes LLPS and forms biomolecular condensates in spinal motor neurons (MNs). Importantly, we discovered that interfering with the K136 SUMOylation site of TDP-43 altered its phase separation behavior, reducing cytoplasmic mislocalization and aggregation. Introduction of the ALS-linked mutation G294V did not alter these LLPS characteristics, indicating that posttranslational modifications such as lysine-specific alterations can modulate TDP-43 pathogenesis through regulating phase separation. Altogether, our *in vivo* characterization of TDP-43 confirms the formation of dynamic nuclear TDP-43 condensates in zebrafish spinal neurons and establishes a critical platform to validate the molecular grammar of phase separation that underpins TDP-43 aggregation in ALS and other proteinopathies.

## INTRODUCTION

For the last 15 years, cytoplasmic TDP-43 aggregates have served as a hallmark feature for several neurodegenerative diseases, including Amyotrophic Lateral sclerosis (ALS) and Fronto-Temporal Dementia (FTD) (Arai et al., 2006; Cairns et al., 2007; Neumann et al., 2006). TDP-43 aggregates are also evident in several other neurodegenerative diseases including Alzheimer’s disease (AD), and limbic predominant age-related TDP-43 encephalopathy (LATE; found in about 25% of individuals above the age of 80 years) (Amador-Ortiz et al., 2007; Huang et al., 2020; Nelson et al., 2019). TDP-43 is a DNA/RNA binding protein (RBP) that is mainly nuclear and plays an important role in RNA metabolism. TDP-43 possesses two RNA recognition motifs (RRMs) allowing the protein to bind RNA and DNA through intronic and 3’UTRs UG-rich regions (Bhardwaj et al., 2013; Buratti and Baralle, 2001; Kuo et al., 2009). TDP-43 interacts with more than 6000 mRNAs, nearly 30% of the entire transcriptome (Tollervey et al., 2011). It plays a major role in RNA processing including transcription, splicing and transport (Lagier-Tourenne et al., 2010). TDP-43’s own RNA expression is regulated by the UG-repeat motifs in its 3’UTR region (Ayala et al., 2011). In addition, TDP-43 can act as a repressor of cryptic exons during RNA splicing, e.g. of *UNC13A*, a gene associated with ALS risk (Brown et al., 2022; van Es et al., 2009; Ma et al., 2021). Overall, the dysregulation of TDP-43 and other RBPs such as FUS, TAF15, ATXN2 or hnRNOA1 is now unequivocally implicated in ALS and other neurodegenerative disease (Xue et al., 2020).

While it is evident that aberrant TDP-43 aggregation and cytoplasmic mislocalization are hallmark features of multiple neurodegenerative diseases, there is still a debate whether degeneration is triggered by loss- or gain-of-function mechanisms (Hergesheimer et al., 2019). TDP-43 may gain new or altered function(s) when it permanently mislocalizes into the cytoplasm (Dammer et al., 2012). Loss of function may result from the nuclear depletion of TDP-43. The importance of nuclear TDP-43 is evidenced by the lethality of animals that lack nuclear TDP-43 (Sephton et al., 2010; Wu et al., 2010). In addition, systematic overexpression of cytoplasmic TDP-43 (ΔNLS TDP-43 variant) also results in accelerated disease phenotypes in mice (Spiller et al., 2016; Walker et al., 2015). These studies highlight the importance of correct compartmentalization for TDP-43 within neurons.

As of today hundreds of post translational modifications (PTMs) have been identified in eukaryotic cells (Khoury et al., 2011). These PTMs have implications for protein localization, stability, solubility, interaction, and degradation and have been known to play an important role in neurodegenerative diseases for many years (Ramani and Zahiri, 2021). Dysregulation of all these processes has been closely linked to pathological inclusions in ALS neurons. PTMs implicated in TDP-43 aggregation and ALS include phosphorylation, acetylation, oxidation, fragmentation, ubiquitination, SUMOylation (Buratti, 2018). For example, in the early 90’s Matsumoto *et al*. found that filamentous inclusions in all ALS patient samples were reactive for ubiquitin (Matsumoto et al., 1990). Ubiquitin positive aggregates became a hallmark feature of the disease before TDP-43 was identified as the main component of aggregates in ALS (Neumann et al., 2006). Furthermore, pathological TDP-43 inclusions are aberrantly phosphorylated and it has been shown that TDP-43 itself is phosphorylated at the double serine 409/410 (Hasegawa et al., 2008; Inukai et al., 2008). Phosphorylation has been associated with toxicity *in vitro* and TDP-43 oligomerization *in vivo* (Choksi et al., 2014; Kim et al., 2015).

SUMOylation is another PTM that was first highlighted in neurodegenerative disorders such as Alzheimer’s and Huntington’s disease (Feligioni et al., 2015) in which SUMOylation has been shown to have a dual effect on aggregation (enhancing or inhibiting), illustrating the importance of well-balanced regulation (Feligioni et al., 2015; Keiten-Schmitz et al., 2021). SUMOylation is a three-step enzymatic process that is known to play a role in nucleo-cytoplasmic transport, protein-protein interaction and stability, mechanisms that are all disrupted in ALS. SUMOylation occurs within a special motif, ψKxE/D, containing the lysine K which can be SUMOylated surrounded by a hydrophobic amino acid (ψ), any residue (x) and an acidic amino acid (either aspartate D or glutamate E). Recently, we have shown that TDP-43 can be SUMOylated by SUMO1 *in vitro* (Maurel et al., 2020). The putative site of SUMOylation was identified as the lysine 136 using bioinformatic tools (Dangoumau et al., 2013). Preventing TDP-43 SUMOylation by mutating its lysine 136 into an arginine, revealed the importance of this residue in TDP-43 cellular localization, cell viability and aggregation (Maurel et al., 2020). Another *in vitro* study recently confirmed the role for the same residue, lysine 136, and SUMOylation, on TDP-43 sub-cellular localization, recruitment to stress granules (SG), as well as splicing activity. The effect of SUMOylation on cellular TDP-43 pathology has so far not been investigated in vivo.

TDP-43 has been shown *in vitro* to undergo liquid-liquid phase separation (LLPS) due to its low complexity domain (LCD) in the C-terminal part of the protein - forming dense liquid-like membrane-less organelles termed biomolecular condensates (BMCs) (Chien et al., 2021; Molliex et al., 2015). PTMs often occur in this LCD domain and have therefore emerged as powerful modulators of LLPS and condensate formation (Hofweber and Dormann, 2019). The concentration of RBPs, such as TDP-43 and FUS, into BMCs is now a well-accepted concept that can help to explain cellular and molecular functions. BMCs can be formed in a short time frame as a result of different stimuli as a protective mechanism or to concentrate biochemical reactions without the need for proteins to be shuttled elsewhere in the cell. The precise mechanisms (e.g. cytoplasmic drivers and molecular kinetics) on how BMCs are modified and regulated in the nucleus or cytoplasm of a cell, are of immense interest as they conceivably provide novel routes to functionally regulate intra- and inter protein interactions (Jord et al., 2021). In neurodegenerative diseases, this concept has generated a “drops become clogs” dogma and emerged as a potential explanation how condensates can phase transition into aggregates (Dolgin, 2018). So far, *in vivo* evidence of this process in TDP-43 regulation has been lacking.

Mutations in the LCD domain of RBDs, including TDP-43, disrupt phase separation leading to fibril formation (Conicella et al., 2016; Molliex et al., 2015). It is therefore important to better understand the molecular mechanisms that drive TDP-43 phase separation mislocalization and aggregation. It is conceivable that the dysregulation of TDP-43 function may be amplified by a liquid-to-solid phase transition (or vice-versa), potentially driving toxic aggregation. The role of PTMs as modulators of LLPS has gained significance over the last years (Hofweber and Dormann, 2019; Loh and Reiter, 2021; Luo et al., 2021). Several *in vitro* studies on other ALS proteins, such as FUS, demonstrated that increased phosphorylation in the LCD domain affected LLPS and aggregation (Monahan et al., 2017; Rhoads et al., 2018). A study from 2016, identified a role of SUMOylation in SG assembly containing eIF4A2, an ATP-dependent RNA helicase (Jongjitwimol et al., 2016). In addition, other studies, reviewed by Keiten-Schmitz *et al*., tried to untangle the role of SUMO in BMCs and highlighted a role in both their assembly and disassembly,. (Keiten-Schmitz et al., 2021).

In this study we aimed to characterize the role of SUMOylation on TDP-43 compartmentalization and phase transition in an *in vivo* zebrafish model. We demonstrate that mutation of the putative SUMOylation site lysine 136 results in enhanced nuclear retention and reduced cytoplasmic aggregation, confirming previous *in vitro* findings. We further provide, to the best of our knowledge, the first characterization of TDP-43 phase separation in spinal MNs of a *in vivo* using the zebrafish model. Comparative analysis of the C-terminal sporadic and familial ALS variant, p.G294V (Del Bo et al., 2009; Williams et al., 2009), revealed that these effect were site specific and not a general phenomenon of mutated TDP-43. Altogether, we present important *in vivo* evidence that TDP-43 accumulates into biomolecular condensates that fuse and separate over time, and that post-translational modifications such as SUMOylation affect TDP-43 phase separation behavior.

## RESULTS

### Lysine 136 influences TDP-43 subcellular compartmentalization *in vivo*

Using bioinformatic tools, lysine 136 (VK_136_KD) has previously been identified as the most likely residue for TDP-43 SUMOylation (Maurel et al., 2020; Zhao et al., 2014). To reveal the impact of lysine 136 *in vivo*, we expressed human variants of eGFP-tagged human TDP-43 in spinal MNs of zebrafish. Driven by a MN specific promoter (*-3mnx1*), we created 4 constructs: the wild-type form of human TDP-43 (TDP-43^WT^), an ALS mutant p.G294V (TDP-43^G294V^), and the SUMOylation-deficient counterparts where the lysine 136 was mutated into an arginine (p.K136R; TDP-43^K136R^ and TDP-43^G294V_K136R^) (fig. 1A). Constructs were sequence-verified, and zebrafish expression was validated by Western Blotting (fig. 1B). Injection of these constructs into the one cell stage of zebrafish eggs lead to mosaic expression, allowing precise characterization of individual eGFP-positive MNs fig. 1C). Injection of these constructs led to mosaic integration, allowing precise characterization of individual eGFP-positive neurons (fig. 1C). First, TDP-43 compartmentalization in MNs was quantified to generate a nucleo-cytoplasmic expression profile of TDP-43 for the different variants. To do so, we used a previously established plot line analysis approach of maximum projection images of GFP positive neurons ((Asakawa et al., 2020); fig. 2A; methods for details). The ratio of fluorescence values (Fmin in the cytoplasm/Fmax in the nucleus) indicates the fluorescence distribution between the cytoplasm and the nucleus, with reduced ratios indicating lower cytoplasmic fluorescence. Both TDP-43^WT^ and TDP-43^G294V^ displayed higher Fmin/Fmax eGFP ratios compared to their SUMO mutated variants (fig. 2B). This change in ratios was mainly driven by reduced cytoplasmic fluorescence intensity as the Fmax fluorescence values within the nucleus and the average intensities (Fmean) along the lines did not change within the assessed variants (Supp fig. 1).

**Figure 1.**
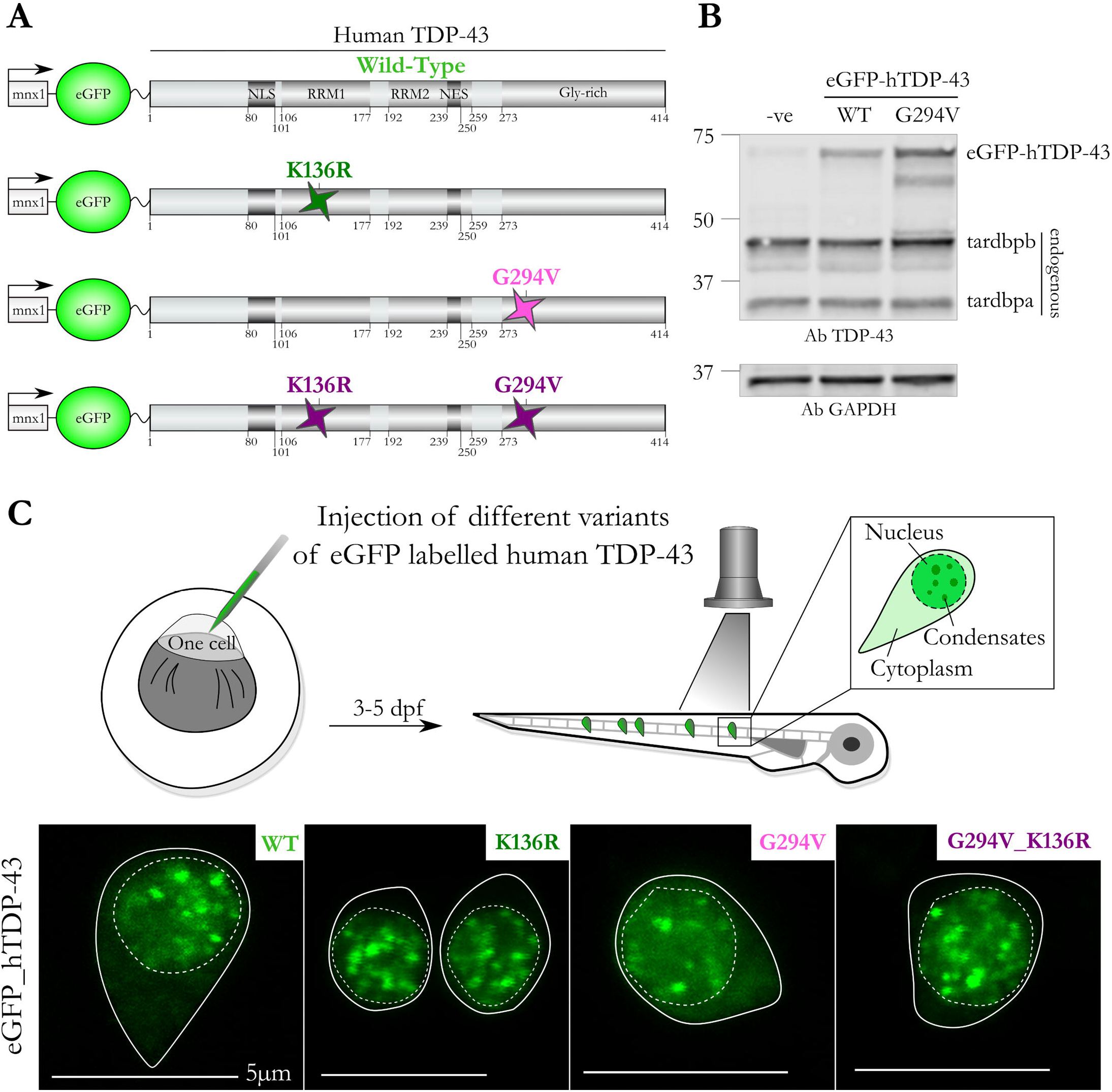
Human TDP-43 expression in the zebrafish spinal cord. (A) TDP-43 constructs with their Nuclear Localization Signal (NLS), two RNA Recognition Motifs (RRMs), Nuclear Export Signal (NES) and glycine–rich domain (low complexity domain). Human TDP-43 was fused to eGFP at the N-terminal and expressed under the MN specific promotor, *-3mnx1*. Four variants were used in this study, wild-type TDP-43^WT^, an ALS variant TDP-43^G294V^, in addition with a single point mutation for the putative SUMOylation site, lysine 136 (TDP-43^K136R^ and TDP-43^G294V_K136R^). (B) Validation of constructs using Western blot analysis of total protein extract of transgenic embryos expressing eGFP-TDP-43^WT^ or ^G294V^ against TDP-43 and GAPDH. Negative siblings were used as a negative control. (C) Constructs were injected in the one cell stage of wild-type zebrafish eggs. Three to five days post fertilization (dpf), embryos expressing GFP positive neurons were imaged using confocal microscopy to characterize eGFP-expression in the cytoplasm (plain line), the nucleus (dotted line), and BMCs. Scale 5 μm.

**Figure 2.**
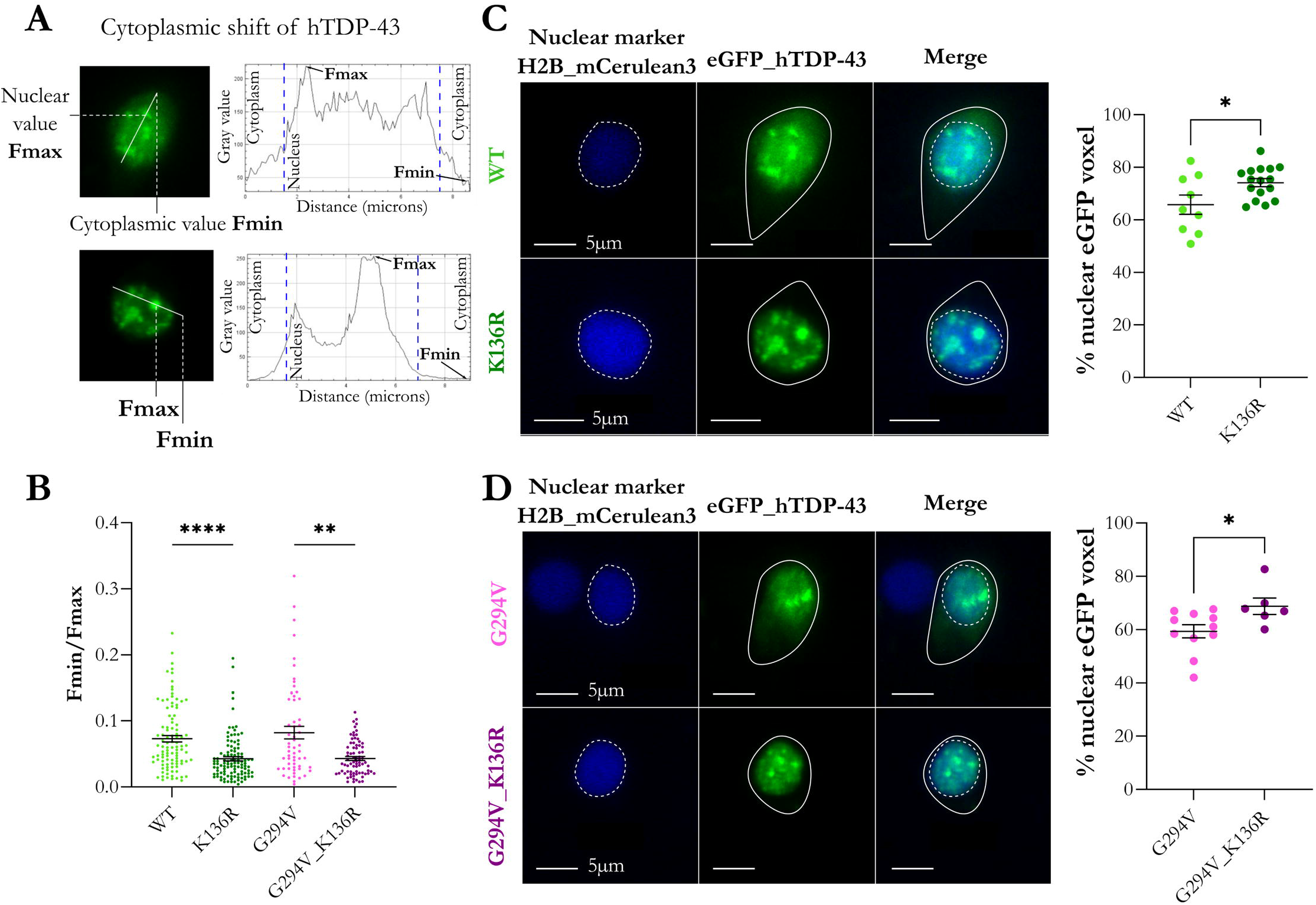
Lysine 136 influences TDP-43 subcellular compartmentalization *in vivo*. **(A)** Plot line profile analysis was used to evaluate the cytoplasmic mislocalization of TDP-43 in MNs by calculating the ratio of Fmin (cytoplasmic) and Fmax (nuclear). Lines were randomly chosen to go through the large part of the nucleus and had to include a droplet. **(B)** Quantitative analysis revealed the importance of lysine 136 in TDP-43 compartmentalization. TDP-43^K136R^ showed a reduced ratio (0.043 ± 0.003) compared to TDP-43^WT^ (0.073 ± 0.004). TDP-43^G294V^ (0.082 ± 0.009) showed the same pattern of a reduced ratio compared to TDP-43^G294V_K136R^ (0.043 ± 0.007). Kruskal-Wallis, n=57-109 neurons in 12-24 larvae, ****p<0.0001 **p=0.0096. **(C-D)** Representative images for TDP-43^WT^ and TDP-43^K136R^ (**C**) and TDP-43^G294V^ and TDP-43^G294V_K136R^ (**D**) expression of nuclear H2B-mCerulean3 (blue) and eGFP-TDP-43 (green). **(C)** 3D volume quantification of GFP distribution in the cell (co-localization with the nuclear marker) revealed a significant higher nuclear TDP-43 retention for TDP43^K136R^ compared to TDP-43^WT^ (74.2% ± 1.5% versus 65.8% ± 3.7%; mean + SEM, unpaired t test, n=9-16 larvae, *p=0.0215). **(D)** TDP-43^G294V^-linked eGFP displayed 59.4% ± 2.74% (mean + SEM) of its GFP signal in the nucleus versus 68.8% ± 3.1% (mean + SEM) for G294V_K136R, n=6-11, unpaired t test, n=6-11, *p=0.0329. Scale = 5 μm.

To verify these observations, we next took advantage of a transgenic zebrafish line expressing the nuclear marker H2B fused to mCerulean3 fluorescent protein (fig. 2C & D). Using a previously published ImageJ macro (Svahn et al., 2018) to determine 3D voxel colocalization of TDP-43, we determined that TDP-43^WT^ displayed 65.8% of its GFP positive voxels colocalized with nuclear mCerulean3. SUMO TDP-43^K136R^ expression showed a significant increase of nuclear GFP-TDP-43 at 74.2% (fig. 2C). These results are in line with previous *in vitro* observations for TDP-43^WT^ (Maurel et al., 2020). Comparison of the lysine 136 mutation in a mutant TDP-43 variant (G294V) revealed a similar outcome. TDP-43^G294V_K136R^ displayed a significantly higher percentage of GFP positive voxels in the nucleus in comparison to the ALS mutant TDP-43^G294V^ (68.8% versus 59.4%) (fig. 2D). In contrast, the ALS mutant TDP43^G294V^ did not show an increased propensity for eGFP-TDP-43 to localize in the cytoplasm compared to the wildtype TDP-43 control.

Altogether, we demonstrate that mutation of the lysine 136 residue in TDP-43 affects its nucleo-cytoplasmic localization and leads to an increased TDP-43 retention within the nucleus of zebrafish MNs. These effects were site-specific and not observed when introducing the C-terminal mutation G294V, found in sporadic and familial ALS patients.

### Cytoplasmic TDP-43 aggregation is affected by an ALS mutation and lysine 136 substitution

We frequently observed in MNs the accumulation of TDP-43 outside the nucleus, representative of cytoplasmic aggregates (fig. 3B), the hallmark feature of TDP-43 pathology (Jucker and Walker, 2013; Neumann et al., 2006). We therefore investigated how these mutations affected the occurrence of such cytoplasmic TDP-43 accumulations (considered aggregates for this analysis) in zebrafish spinal cord neurons. 18.3% of MNs expressing TDP-43^WT^ displayed such cytoplasmic aggregates compared to 26.4% of the MNs expressing the disease-associated TDP43^G294V^ mutation (fig. 3A). Notably, the SUMOylation-deficient counterparts (TDP43^K136R^ and TDP43^G294V_K136R^) displayed a significantly reduced percentage of cells with these aggregates (13.2% and 4.2% respectively, fig. 3A). The number of aggregates within the cytoplasm of individual neurons was also reduced for the SUMO-deficient variants (TDP43^K136R^ and TDP43^G294V_K136R^) compared to the WT and p.G294V variants (fig. 3B). Altogether, these data demonstrate the capacity of the lysine 136 residue to affect the cytoplasmic accumulation of TDP-43-while retaining TDP-43 in the nucleus.

**Figure 3.**
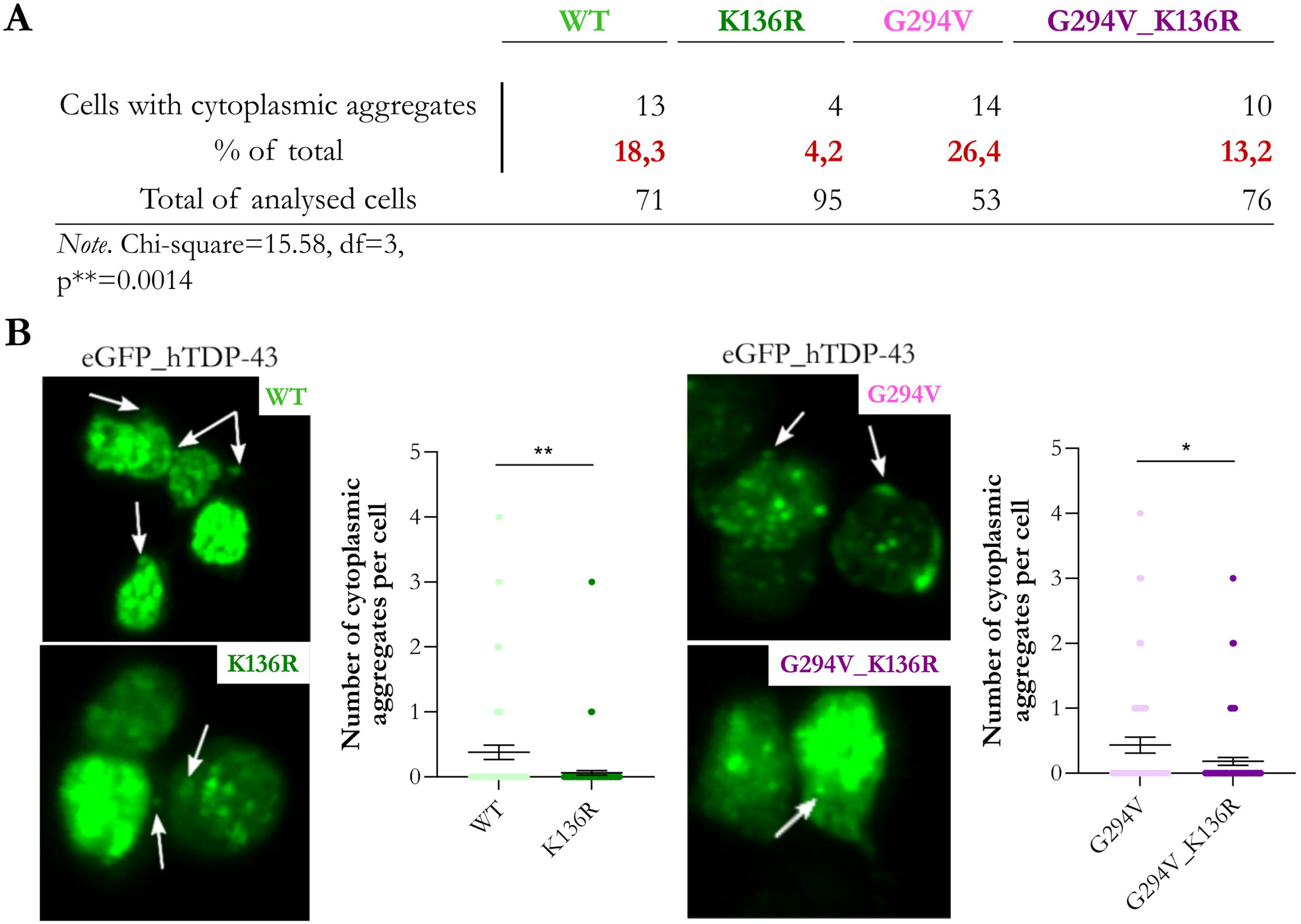
Cytoplasmic TDP-43 aggregation is affected by an ALS mutation and lysine 136 substitution. **(A)** Quantification of cells positive for eGFP-HsaTDP-43 presenting cytoplasmic accumulation (representative of cytoplasmic aggregation here) of TDP-43. Zebrafish spinal MNs with the ALS mutation G294V displayed higher percentage of cytoplasmic aggregation. Addition of the K136R mutation resulted in a decrease of cytoplasmic aggregation (n_WT_=71, n_G294V_=53, n_K136R_=95, n_G294V_K136R_=76, Chi-square test **p=0.0014). **(B)** Quantification of the cytoplasmic aggregates per cell showed a reduced number of aggregates for both SUMO-TDP-43 variants. Left panel: TDP-43^WT^ and TDP-43^K136R^ (0.38 ± 0.11 versus 0.06 ± 0.03 respectively; Mann-Whitney test; **p=0.0017). Right panel: TDP-43^G294V^ and TDP-43^G294V_K136R^ (0.43 ± 0.12 compared to 0.18 ± 0.06 cytoplasmic aggregates; Mann-Whitney test *p=0.0484).

### Lysine 136 affects characteristics and dynamics of nuclear TDP-43 condensates *in vivo*

We previously reported the accumulation of highly dynamic TDP-43 droplets in the nucleus of MNs expressing TDP-43^WT^ (Svahn et al., 2018). Over the last years, phase separation has emerged as a novel concept responsible for the formation of such accumulations. More precisely, *in vitro* studies identified a mechanism, called liquid-liquid phase separation (LLPS), to prompt the segregation of proteins such as TDP-43 into two distinct liquid phases, i.e., a diffuse phase, and a highly condensated phase, within the same physiological environment. Those biomolecular condensates (BMCs) can mix and de-mix upon different stimuli, allowing rapid rearrangement and diffusion of RNA and proteins into and out the condensate (fig. 4A). It is now understood that mis-regulation of TDP-43’s BMC formation through LLPS can lead to liquid-to-solid phase separation (LSPS) and protein aggregation in neurodegenerative diseases such as ALS (Maharana et al., 2018; Prasad et al., 2019; Ramaswami et al., 2013). We therefore explored whether human TDP-43 expressed in our zebrafish displayed such a phase separation behavior.

**Figure 4.**
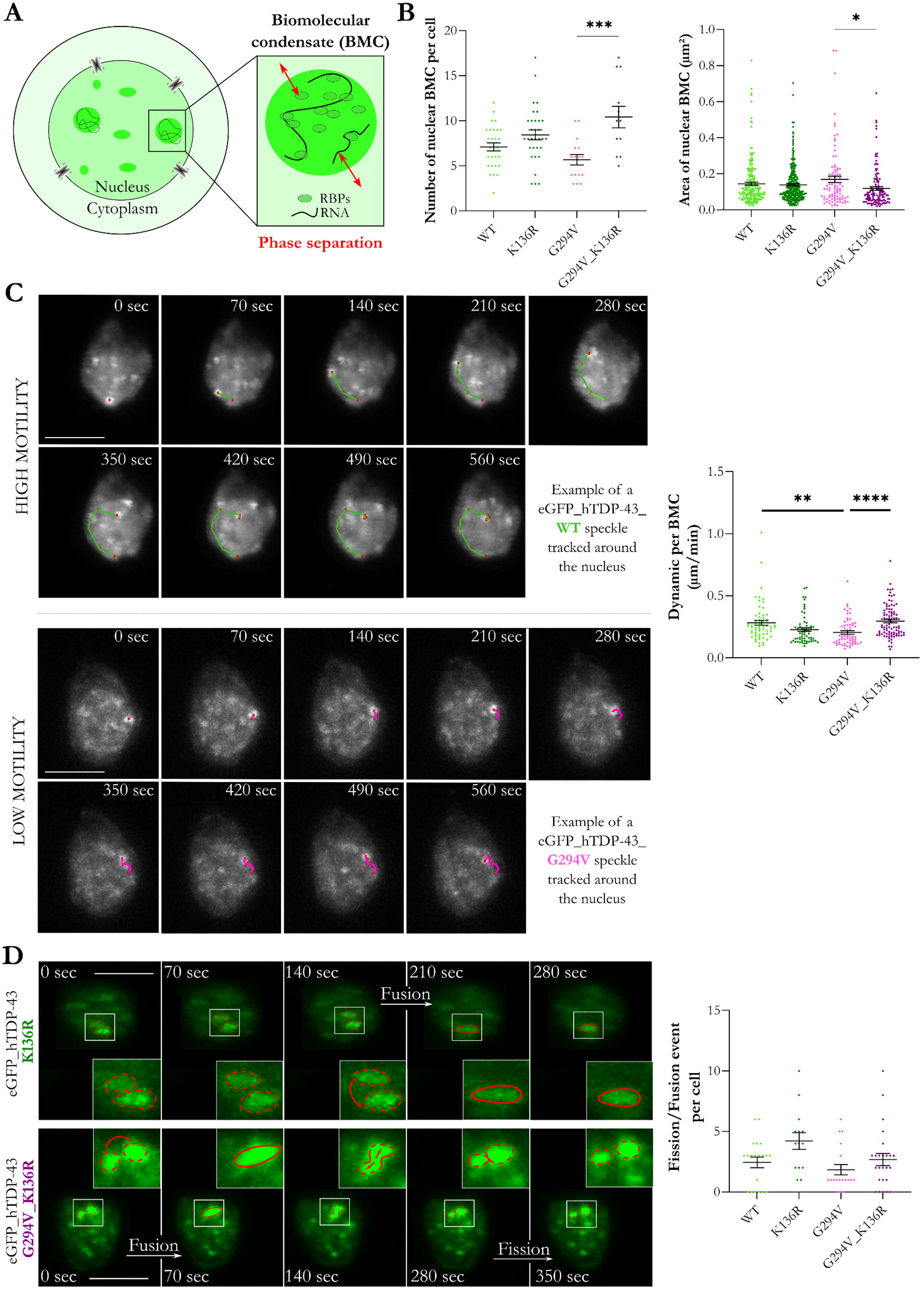
TDP-43 forms condensates within the nucleus of MNs *in vivo* and mutation of lysine 136 affects these condensate characteristics and dynamics. (A) Schematic illustrating condensates containing RNA (black line) and RNA binding proteins (such as TDP-43) in the nucleus and undergoing phase separation with a continuous exchange within the nuclear environment (red arrows). (B) Quantification of BMC numbers per MN (7.1 ± 0.5 BMC per cell for TDP-43^WT^, 8.4 ± 0.6 BMC per cell for TDP-43^K136R^, 7.1 ± 0.5 BMC per cell for TDP-43^G294V^ and 8.4 ± 0.6 BMC per cell for TDP-43^G294V_K136R^, One-Way ANOVA test, ***p=0.0004). Quantification of the area of those BMCs (0.145 μm^2^ ± 0.009 for WT, 0.139 μm^2^ ± 0.006 for K136R, 0.170 μm2 ± 0.018 for G294V and 0.118 μm2 ± 0.010 for G294V_K136R, Kruskal-Wallis test, n=91-270 speckles in 12-32 neurons, *p=0.0197). (C) Representative images showing the trajectory of individual droplets inside the nucleus (maximum projection images). Top panel shows a highly motile condensate (TDP-43^WT^) and the bottom panel represents a low motility condensate (TDP-43^G294V^). Dynamics data were registered every 70 seconds for a maximum of 21 minutes (0.282 μm/min ± 0.019 for WT, 0.204 μm/min ± 0.014 for G294V, 0.227 μm/min ± 0.014 for K136R, 0.296 μm/min ± 0.014 for G294V_K136R, Kruskal-Wallis test, n=62-92 tracking in 11-14 neurons, **p=0.0010 ****p<0.0001). (D) Representative examples and quantification of fission and fission events characterizing LLPS and BMC in the nucleus (2.5 events ± 0.4 for WT, 1.8 events ± 0.4 for G294V, 4.2 events ± 0.7 for K136R, 2.7 events ± 0.5 for G294V_K136R, Kruskal-Wallis test, n=14-25, ns). Scale 5 μm.

First, we compared the number of nuclear *in vivo* droplets. We observed no differences in the number or area of droplets between TDP-43^WT^, TDP-43^K136R^ and TDP-43^G294V^, except in the TDP-43^K136R_G294V^ double mutation where we found an increased number of droplets which were also significantly reduced in area (fig. 4B). We next assessed the dynamics by tracking individual TDP-43 droplets within the nucleus using time-lapse confocal imaging (Supp. Video 1). Maximum projection images (MIPs) were used to track unequivocally identified and individual droplets over ~10 minutes. The TDP-43^G294V^ mutant displayed a lower motility compared to TDP-43^WT^, covering shorter distances over the same time period. Figure 4C illustrates two representative examples of these different motilities for both a TDP-43^WT^ and TDP-43^G294V^ positive MN (see Supp. Video 1 for the timelapse video). While TDP-43^K136R^ showed a slightly reduced motility compared to the WT form of TDP-43, this change was not significant. Introducing the p.K136R mutation in the p.G294V variant however restored the motility of the droplets to a level comparable to TDP-43^WT^ (fig. 4C).

Another hallmark feature of BMCs is their ability to fuse and separate over time. Indeed, we observed spontaneous condensate fusion and fission over minutes within the nucleus in zebrafish MNs (fig. 4D and Supp. Video 2). Comparing the number of fusion and fission events of the different TDP-43 variants revealed an increased tendency of the SUMO-deficient variants to undergo these events (not statistically significant) (fig. 4D).

Altogether, we provide the first detailed *in vivo* cellular characterization of TDP-43 nucleation, condensate dynamics, and aggregation in zebrafish spinal MNs. While SUMO-deficient variant of TDP-43^K136R^ showed limited changes compared to the WT form, TDP-43^G294V_K136R^ led to significant changes in droplet number, area and motility compared to its TDP-43^G294V^ counterpart.

### Lysine 136 influences the molecular dynamics of TDP-43 and biomolecular condensate (BMC) formation

Fluorescence Recovery After Photobleaching (FRAP) is technique widely used to determine the molecular characteristics of BMCs and the exchange of proteins within their environment (Kuzma-Kuzniarska et al., 2016; Vink et al., 2006). We therefore established a FRAP approach to measure the molecular characteristics and exchange of TDP-43 within these BMCs in our spinal MNs *in vivo* (see methods for details).

Using a short UV laser pulse, we bleached one single droplet within the nucleus of a MN, quenching the existing GFP signal. Upon sufficient bleaching (>50%; see methods), we measured the recovery of the eGFP signal over time (fig. 5A). This recovery was relatively quick, around 60 seconds to restore 50% of the initial baseline signal for TDP-43^WT^. The normalized data were curve fitted using a double term exponential equation (*I*_*fit*2_ = *I*_0_ – *α.e*^−*β.t*^ – *γ.e*^−*δ.t*^) to allow the calculation of recovery times and the mobile and immobile fractions of TDP-43 (Supp fig. 2). T-half time represents the half maximal recovery time (the speed of recovery), and the mobile fraction determines the percentage of molecules that are free to diffuse back into droplets. Comparing WT to p.G294V revealed no differences for FRAP recovery half-time and mobile fractions (fig. 5B & C). However, assessing the recovery curves of the SUMO TDP-43 variants revealed significant recovery differences. Fluorescence recovery was strikingly faster when lysine 136 was additionally mutated for both variants compared to WT (T-half time of 59.3 sec for WT; 13.1 sec for K136R) and p.G294V (T-half time of 130.5 sec for G294V; 15.7 for G294V_K136R) (fig. 5B & C). The mobile fraction was increased for all TDP-43 variants compared to TDP-43^WT^ revealing a significant difference for TDP-43^K136R^ (fig. 5C). We questioned whether the amount of diffuse TDP-43 in the nucleus, compared to the bound TDP-43 within the BMCs, was altered between the variants and could contribute to the observed difference in FRAP recoveries (fig. 5D). Indeed, the fluorescence intensity ratios of diffuse TDP-43 in the nuclear background (without BMCs) to specific condensate-bounded TDP-43 fluorescence were significantly reduced when lysine 136 was mutated (fig. 5D), indicating an increased nucleation propensity of (diffuse) TDP-43 into BMCs within the nucleus in the SUMO-deficient variants TDP-43^K136R^ and TDP-43^G294V_K136R^. Fluorescence intensities of individual BMCs and the diffuse TDP-43-eGFP background were not altered (Supp fig. 3). Nuclear fluorescence plot profile analysis of MIPs confirmed this observation that mutation of lysine 136 only reduces the amount of diffuse TDP-43 within the nucleus (Supp fig. 4).

**Figure 5.**
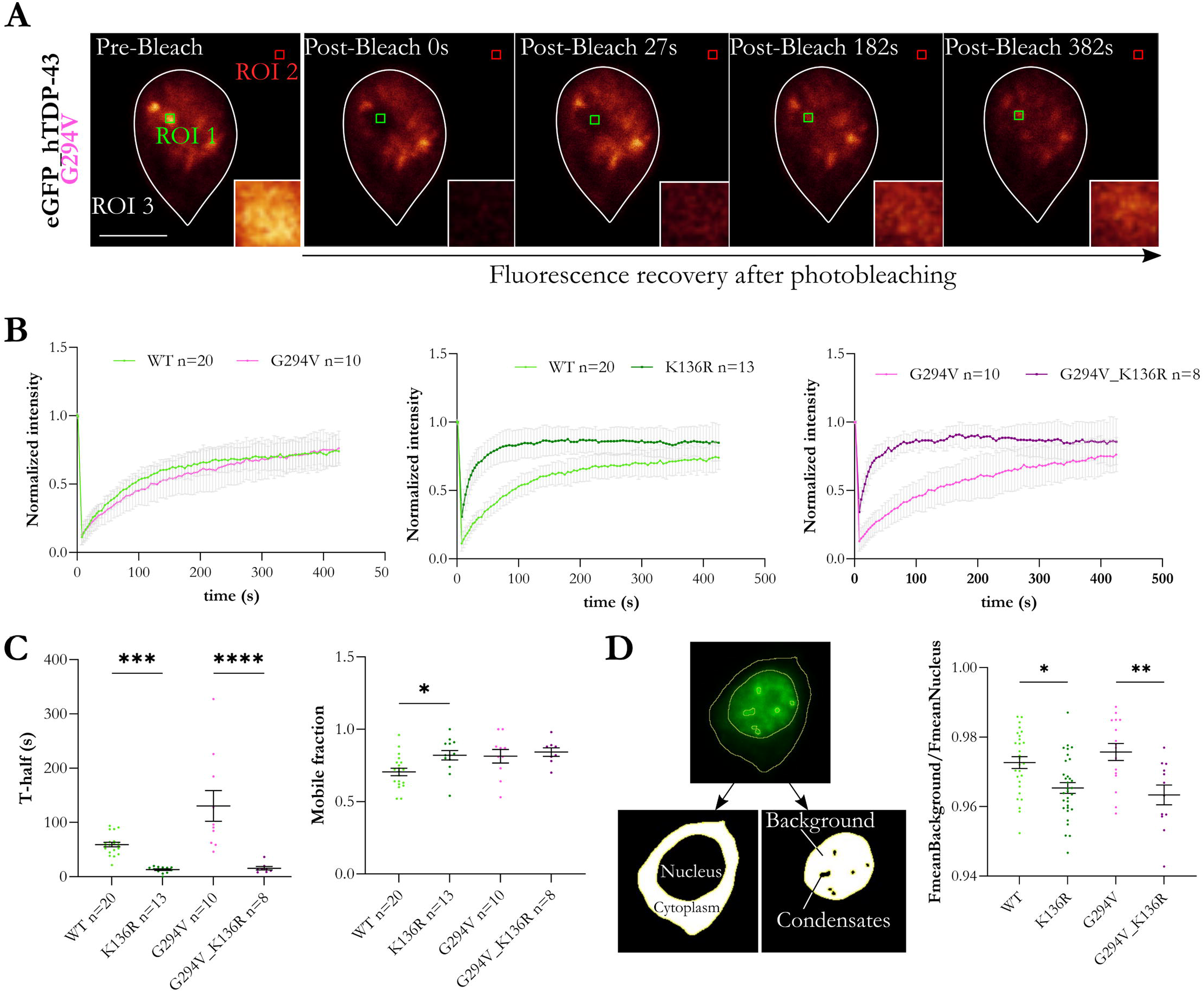
Lysine 136 influences the molecular dynamics of TDP-43 and biomolecular condensate phase characteristics. **(A)** Representative images illustrating Fluorescence Recovery After Photobleaching (FRAP) in spinal MNs *in vivo*. Using UV, the region of interest (ROI1, the BMC) was photobleached and the recovery of fluorescence was measured afterwards. ROI2 (background) and ROI3 (reference, total fluorescence) were used to normalize FRAP recovery. **(B)** Average normalized FRAP curves for all tested conditions. **(C)** Scatter plots comparing T-half time and mobile fraction extracted from double term exponential equation fitted curves (T-half time of 59.3 seconds ± 4.3 for WT, 130.5 seconds ± 28.4 for G294V, 13.1 seconds ± 1.6 for K136R, 15.7 seconds ± 3.2 for G294V_K136R, Kruskal-Wallis test, n=8-20, ***p=0.0001 ****p<0.0001 // mobile fraction of 0.71 ± 0.03 for WT, 0.81 ± 0.05 for G294V, 0.82 ± 0.03 for K136R, 0.84 ± 0.03 for G294V_K136R, One-way ANOVA test, n=8-20, *p=0.041). **(D)** Analysis of diffuse TDP-43 in the background (without BMCs) compared to the overall fluorescence of the nucleus (ratio FmeanBackground/FmeanNucleus of 0.973 ± 0.002 for WT, 0.976 ± 0.002 for G294V, 0.965 ± 0.002 for K136R, 0.963 ± 0.003 for G294V_K136R, One-way ANOVA test, n=12-32, *p=0.0151 **p=0.0038).

To the best of our knowledge, this is the first *in vivo* demonstration of TDP-43 phase separation using FRAP techniques in the zebrafish spinal cord. Analysis revealed a strong effect of lysine 136 on TDP-43 phase separation, supporting the idea that PTMs can directly affect LLPS behavior.

## DISCUSSION

In this study we investigated the role of SUMOylation, more precisely lysine 136, on TDP-43 dynamics, liquid-liquid phase separation (LLPS), and neuronal compartmentalization *in vivo*. Our zebrafish findings emphasize the distinct effects that a single lysine mutation on TDP-43 can have on nuclear localization and cytoplasmic aggregation. We demonstrate that TDP-43 assembles in spinal MNs into distinct biomolecular condensates (BMCs) through LLPS. Modification of lysine 136 revealed reduced cytoplasmic expression, decreased aggregation, and accelerated molecular exchange compared to wild-type and disease-associated TDP-43 (p.G294V). Our results highlight the importance of molecular drivers such as post-translational modifications, mutations, and potential interactors for phase transition and condensate formation as well as aggregation in neurodegeneration *in vivo*. Our novel workflow further closes a critical gap in the field to analyze condensate formation and phase separation at a single cell level and in a living organism.

TDP-43 is an RNA binding protein (RBP) that is predominantly located within the nucleus. TDP-43 shuttles in and out the nucleus and has been demonstrated to play important roles in RNA metabolism (Weskamp and Barmada, 2018). In ALS and other neurodegenerative disease, this well-regulated process is disrupted, subsequently leading to the formation of cytoplasmic inclusions as the hallmark feature for ALS (~97% of patients) and FTLD (~50% of patients). TDP-43 contains two RNA-recognition motifs (RRM1 and 2) and a C-terminal low complexity region (LCD), also referred as an intrinsically disordered domain (IDR), which both play important roles in LLPS (Babinchak et al., 2019; Conicella et al., 2016). Generally, strong cation-π interactions that involve aromatic residues (such as phenylalanine [F], threonine [T], tryptophane [W]) and positively charged residues (such as arginine [R], lysine [K], asparagine [N], glutamine [Q], or tyrosine [Y]) are considered important drivers for phase separation (Wang et al., 2018b). RGG/RG motifs have also been identified as powerful modulators of LLPS, generally through RNA-binding interactions (Chong et al., 2018). Interestingly, TDP-43 contains one RGG motif in its LCD (Supp fig. 5). Methylation of TDP-43’s arginine at the position 293 has been shown to regulate liquid-to-solid phase transition (Aikio et al., 2021), and glycine residues have also been identified to affect condensation (Wang et al., 2018b). Out of the 141 residues composing TDP-43’s LCD, 38 are glycine (C-terminal), 22 are serine and 11 glutamine residues (with many reported to be altered by PTMs or mutated in MND). However, the precise effects on how specific amino acids influence TDP-43 condensation and aggregation are just beginning to emerge and are likely not limited to the IDR or the RNA-binding domains.

For instance, the single phosphomimic substitution (an amino acid modification to replicate a phosphorylated residue by changing serine to glutamic (E) or aspartic acid (D)) of the S48 by a glutamic acid (E) disrupts phase separation as well as TDP-43’s N-terminal capacity to self-oligomerize (Wang et al., 2018a). Glycine 338 located in the α-helical structure of the LCD of TDP-43’s also influenced FRAP recovery following site-directed mutagenesis to alanine *in vitro* using HeLa cells (Conicella et al., 2020). Cohen et al. revealed the importance of the acetylated lysines (145 and 192, respectively in the RRM1 and RRM2 domain of TDP-43) in TDP-43 insolubility and its aggregation propensity (Cohen et al., 2015). Thus, there is mounting evidence that single point mutations can strongly influence TDP-43’s functional properties, conceivable through altering LLPS propensities. Furthermore, a recent study investigating the impact of phosphomimic substitutions in TDP-43 (serine to aspartic acid) reported that hyperphosphorylation affected LLPS and might be protective under specific circumstances (Silva et al., 2021). The authors demonstrated that an increased number of phosphomimic mutations (12D compared to 5D and WT-TDP-43) reduced FRAP recovery rates, increased BMCs dynamics, and reduced TDP-43 aggregation. Our *in vivo* SUMOylation results align with these observations that PTMs can have protective effects on protein aggregation and mislocalization, likely through influencing phase separation.

Lysines have been shown to play an important role in aggregation and phase separation of proteins *in vitro* (e.g. tau, Ukmar-Godec et al., 2019). Lysines are subjected to many PTMs (e.g. ubiquitination, methylation, acetylation and SUMOylation) that can impact physiological function and cellular processes (Wang and Cole, 2020). For instance, TDP-43’s lysine 136 is prone to acetylation and Garcia Morato *et al*. showed that acetyl-mimic K136Q reduces RNA binding and that both acetyl-mimic and acetyl-dead K136R disrupts exon 9 *CFTR* splicing activity, implicating RNA interactions in LLPS (Garcia Morato et al., 2022). Further analysis of K136Q condensates revealed reduced motility, size and fusion events compared to TDP-43^WT^. Importantly, the authors showed that deacetylation (via sirtuins) reduced TDP-43 acetylation and resulted in a reduction of aggregation *in vitro*. The same 136 lysine is also predicted to be a strong candidate for SUMOylation and we have recently shown *in vitro* that TDP-43 is indeed SUMOylated (Maurel et al., 2020). TDP-43 is an RNA binding protein (RBP) that is predominantly located within the nucleus, and we have reported that the SUMO-deficient K136R version of TDP-43 affects its cellular localization, with an increased nuclear retention compared to the WT variant. These findings for SUMOylation of TDP-43 nucleocytoplasmic distribution *in vitro* have since been confirmed (Maraschi et al., 2021), and SUMOylation has also been shown to promote LLPS of Sop-2, an epigenetic regulator of transcription, in *C. elegans* (Qu et al., 2020). However, to the best of our knowledge, phase separation of TDP-43 has never been assessed in a living vertebrate before. Our *in vivo* results confirm that TDP-43 is highly motile and forms condensates predominantly in the nucleus, which can undergo spontaneous fusion and fission events. These characteristics stipulate that TDP-43 forms BMCs in spinal zebrafish MNs. FRAP analysis revealed a striking shift in the fluorescence recoveries in p.K136R condensates compared to controls, conceivably indicating an increase of the molecular dynamics of these BMCs. Remarkably, these effects of aberrant phase separation were limited to the mutation of the single lysine 136 while the disease-linked mutation (G294V) had no effect on these LLPS characteristics. Garcia and colleagues reported in their *in vitro* study that the acetylation-mimic variant K136Q (instead of acetylation- and SUMO-deficient K136R) showed a slower FRAP recovery and formation of larger aggregates *in vitro* (Garcia Morato et al., 2022). These findings are aligned with our *in vivo* LLPS observations of the K136R variant showing the opposite effects. However, considering lysine 136 on TDP-43 can undergo either acetylation or SUMOylation, the shifts that we observed in FRAP recoveries cannot be linked unequivocally to SUMOylation deficiency – and studying SUMO-mimic TDP-43 (instead of SUMO-deficient TDP-43) is experimentally more complex as SUMOylation correspond to the addition of a canonical consensus motif (ψKxE/D) via a 3 steps enzymatic pathway. Nonetheless, using SUMOylation prediction software (http://sumosp.biocuckoo.org/online.php), lysine 136 is identified to be the most preferential residue to be SUMOylated. Further untangling the precise contributions of PTMs such as acetylation and SUMOylation will provide important new knowledge that may help develop pharmacological strategies to moderate TDP-43 pathogenesis in the future.

A limitation of our approach is the overexpression of TDP-43 in our zebrafish spinal MNs. It is evident that BMC dynamics are linked to protein concentration, pH or temperature, and that alterations of such parameters can directly influence LLPS (Zeigler et al., 2021). However, overexpression *per se* in our zebrafish does not seem to be the determining factor, as the disease linked mutation G294V did not reveal aberrant LLPS dynamics compared to WT TDP-43. Additionally, our fish overexpress GFP-tagged human TDP-43 instead of endogenous zebrafish TDP-43, and zebrafish and human TDP-43 show only 57% sequence homology, while murine TDP-43 has 96% homology with the human protein. However, we cannot explicitly untangle the potential influence of endogenous TDP-43 on our observations. While our zTDP-43 expression levels were not altered when hTDP-43 was overexpressed (see Fig. 1), future CRISPR studies that either knock-out endogenous zTDP-43 or introduce small fluorescent tags on endogenous TDP-43 will be important to further delineate the role of LLPS in protein aggregation. Altogether, our zebrafish model provides a novel and powerful platform to investigate how mutations, PTMs, LLPS, and RBPs are contributing to the complex interplay in protein aggregation and consequently disease pathology. So far, our understanding of phase separation has mainly emerged from *in vitro* studies, in simplified model systems as well as more complex living cells. However, many questions remained around the relevance of the biochemical behavior of condensates in more complex living systems where condensate composition and chemical environments are likely to be distinct (Kilgore and Young, 2022). In our fish, we are able to (i) express human TDP-43 in MNs specifically, (ii) assess dynamic phenomena at a subcellular level, and (iii) evaluate distinct molecular changes to investigate molecular mechanisms such as LLPS *in vivo*. With TDP-43 aggregation being found in other diseases, such as Alzheimer’s disease or LATE, this platform provides a critical addition for investigations of *in vivo* phase separation as well as neurodegeneration in general and can be directly applied to other proteinopathies.

Considering aberrant LLPS seems to be critical for the conversion of liquid-like condensates to a solid/gel like aggregates, identification of the molecular signature of LLPS is crucial to decipher the underlying mechanisms that can cause pathological aggregation (Carey and Guo, 2022). For example, LLPS perturbations of ALS mutated FUS droplets lead to fibrous accumulations, synonymous of solidification and pathological aggregation (Patel et al., 2015). Similar fibrillization has been demonstrated for another RBP relevant in ALS, hnRNPA1 (Molliex et al., 2015). Altering the factors (such as PTMs) that are affecting LLPS and transitioning proteins towards a solid like state, could provide novel opportunities to alleviate disease burden (e.g., sirtuins that impact on acetylation level of TDP-43). Small molecule inhibitors are an emerging therapeutic option by targeting aggregation pathways, including by modulating LLPS. For example, IGS-2.7, a casein kinase-1A inhibitor of TDP-43 phosphorylation, has shown reduce cytoplasmic mislocalization of TDP-43 in lymphoblasts from ALS patients. Moreover, it increased the viability of MNs in an ALS mouse model (Martínez-González et al., 2020). Additionally, using a PARylation inhibitor, Olaparib, Duan and colleagues showed a reduced cytotoxicity of TDP-43 *in vitro* (Duan et al., 2019). PARylation is known to enhance LLPS with TDP-43 possessing a PAR-binding motif in the N-terminal domain. However, whether Olaparib directly influences LLPS is still unknown. Also, new emerging molecules have been described *in vitro* to modulate LLPS via stress granules formation. For example, trimethylamine N-oxide (TMAO), a chaperone molecule, can boost TDP-43 condensation, but reduces protein fibrillation *in vitro* (Choi et al., 2018).

Unravelling the exact mechanisms and the molecular and chemical grammar of aggregate formation will continue to have significant impact on the field and help to develop novel therapeutic strategies for modulation of protein aggregation in MND/ALS and other proteinopathies. Especially with the emerging evidence that PTMs may have protective effects or that small-molecule drugs can selectively accumulate into specific condensates.

## METHODS

### Zebrafish studies

Experiments were conducted using zebrafish (*Danio rerio*) under Macquarie University Animal Ethics and Biosafety approvals (2015/033; 52019045211002). All zebrafish were maintained under standard conditions on a 14:10 light:dark cycle with twice daily feeding of artemia and standard pellet at 28 °C (Westerfield, 2000). Larvae were raised in E3 medium (5 mM NaCl, 0.17 mM KCl, 0.33 mM CaCl_2_, and 0.33 mM MgSO_4_ buffered to 7.3 pH using carbonate hardness generator (Aquasonic), no methylene blue). Experiments were conducted at 3–5 days post-fertilization (dpf) unless otherwise stated and under anesthesia using 0.01% (w/v) tricaine methansulfonate (MS-222, Sigma).

### Constructs

Transgenes were generated using MultiStie Gateway® Three-Fragment Vector Construction Kit into recombined pTol2pA2 vector (Kwan et al., 2007). *Tg(mnx1:GAL4,UASEb1:H2B-mCerulean3-P2A-KOFP2-CAAX*) was produced with p5E--3mnx1 (a motoneuron specific promotor neuron) (Morsch et al., 2015), pME-GAL4:UASEb1, p3E-H2B-mCerulean3-P2A-KOFP2-CAAX entries (Don et al., 2016) as described in (Svahn et al., 2018). The nuclear H2B marker is fused to mCerulean3 fluorophore, the membrane localized CAAX marker is fused to mKOFP2 (mKO2) and allows the two proteins to separate. The codon-optimized H2B–mCerulean3–P2A–mKOFP2–CAAX sequence was ordered from GeneArt and recombined into pDONRP2R-P3 (Invitrogen) to create the p3E-H2B-mCerulean3-P2A-mKOFP2-CAAX (Svahn et al., 2018). This transgenic line has been established for nucleo-cytoplasmic studies (Svahn et al., 2018).

*Tg(-3mnx1:eGFP-HsaTDP-43WT*) was generated using recombined p5E-3mnx1 (Morsch et al., 2015), pME-EGFP (Kwan et al., 2007), p3E-HsaTDP-43 and pTol2pA2 (Kwan et al., 2007). The p3E-HsaTDP-43 was generated by subcloning a BamHI-Spe1 flanked genestring encoding HsaTDP-43 (GeneArt) into the BamHI-Spe1 sites of p3E-MCS (Don et al., 2016). SUMO deficient TDP-43 variants TDP-43^K136R^ and TDP-43^G294V_K136R^ were generated by GeneScript mutation services altering c.405-407CAA>CAG and c.405-407CAA>CAG_c.880 GGG>GTT nucleotide sequences respectively.

### Western Blotting

Transgenic lines were generated by co-injection of the Tol2-flanked constructs and transposase mRNA into one-cell stage of zebrafish lines as previously reported (Don et al., 2016; Kawakami, 2004; Morsch et al., 2015; Svahn et al., 2018). 3 to 6 dpf embryos were euthanized and collected (n=10-30) and washed with Ringer Solution (58.2mM NaCl 4.0mM KCl 4.8mM NaHCO3 pH 7) and PBS before lysis in RIPA buffer (50mM Tris-HCl pH 7.4, 150mM NaCL, 1mM EDTA, 1% Triton-X-100, 1% NaH2PO_4_, 0.1% SDS) with cOmplete™, EDTA-free Protease Inhibitor Cocktail and PhosSTOP™ Phosphatase Inhibitor Cocktail (Roche). Proteins were quantified by BCA protein assay, separated by 4-15% SDS-PAGE (BioRad), transferred and treated with a polyclonal rabbit anti-TDP-43 antibody (ProteinTech; 10782-8-AP; 1/10000) and a mouse monoclonal anti-GAPDH antibody (ProteinTech, 60004-1-Ig, 1/20000) overnight at 4°C. Li-Cor secondary antibodies donkey anti-rabbit 680LT and donkey anti-mouse 800CW were used for revelation on Li-Cor Odyssey CLx (Li-Cor, LCR-926-68023 and LCR-926-32212 respectively, 1/15000).

### Imaging

Imaging was conducted on a Leica SP5 or SP8 confocal microscope as previously described (Formella et al., 2018; Morsch et al., 2015, 2017). mCerulean3 and eGFP were excited by 405 nm and 488 nm respectively. High resolution images were obtained on the SP8 confocal microscope while the SP5 confocal microscope was used for FRAP analysis (see below). Objectives used were a Leica 40x/0.80 HCX APO L U-V-I water immersion on the SP5 and HC PL APO CS2 40x/1.10 water immersion on the SP8.

### Nucleo-Cytoplasmic ratio analysis

#### 3D volume analysis of TDP-43 nuclear/cytoplasm ratio

The ratio of nuclear to cytoplasmic eGFP-TDP43 (WT or mutated) over the 3D neuron volume was derived by first establishing a separation and quantification of voxels previously reported (Svahn et al., 2018). One mask contains eGFP-TDP43 (WT or mutated) alone (cytoplasm) or both eGFP-TDP43 (WT or mutated) and H2B-mCerulean3 (nucleus) using a custom macro in ImageJ. The comparison of fluorescence grey value of each voxel sums up of each mask indicated the fluorescence concentration. Single neuron were imaged using a Leica SP8 confocal microscope with identical microscope excitation and collection settings. Multiple technical replicates have been performed to reach the n numbers presented in the study using the same *Tg(mnx1:GAL4,UASEb1:H2B-mCerulean3-P2A-KOFP2-CAAX*) line.

#### Plot line profile

The plot line profiles of TDP-43 (WT or mutated) were assessed by drawing a line through the nucleus (including BMC) and plotting the fluorescence grey values along the y axis. Plot profiles were established using maximum intensity projections (ImageJ) of eGFP-HsaTDP-43 (WT or mutant) positive neurons in the spinal cord of non-transgenic fish (WT TAB). The highest value was determined as Fmax (generally a BMC in the nucleus), and the lowest value as Fmin (generally in the cytoplasm). Cytoplasmic compartmentalization of TDP-43 was evaluated as a relative value of minimal (Fmin, cytoplasm) and maximal (Fmax, nuclear) fluorescence intensity. Lower Fmin/Fmax ratios reflect higher accumulation of eGFP-TDP43 in the nucleus or reduced expression in the cytoplasm. The neurons used in this analysis were imaged with identical microscope excitation and collection settings.

### Cell segmentation and fluorescence analysis

To determine nuclear, cytoplasmic, and BMC fluorescence intensities, maximum projection images of eGFP-HsaTDP-43 (WT or mutant) positive MNs in the spinal cord of non-transgenic fish (WT TAB) were analyzed using a custom macro in ImageJ. Masks were created using non-subjective thresholds (‘Otsu dark’ for the nucleus, ‘Find Maxima’ for the BMCs, and ‘MinError dark’ for the cytoplasm) and all MNs were analyzed using the same settings. Masks were visually confirmed and area and fluorescence grey values were used for analysis. The mean of fluorescence grey values in the nucleus and background were used to calculate the intensity of diffuse TDP-43 in Figure 5D. The FmeanBackground (diffuse TDP-43 without BMCs) was normalized to FmeanNucleus (all nuclear signal with both diffuse and condensated TDP-43) scores.

### BMC movement analysis

Condensate dynamics were evaluated on 2D maximum intensity projections of eGFP-HsaTDP-43 (WT or mutant) positive neurons in the spinal cord of non-transgenic fish (WT TAB). Images were taken every 70 seconds with a 600Hz bidirectional scan speed, with a maximum of 18 timepoints, averaging around 21 minutes per session. Global movement of the cell during imaging was corrected with StackReg (RigidBody) plugin in ImageJ. The diameter and area of the analyzed condensate were collected. Several condensates per cell were tracked when possible, using the Manual Tracking plugin. The condensate’s center was located and tracked on each time-lapse image on 2D maximum intensity projection (MIP). The overall distance was divided by the number of timepoints (μm per 70 seconds) and followed calculation to express in μm per minute.

### FRAP analysis

Fluorescence recovery after photobleaching was performed on the Leica SP5 confocal microscope using the FRAP wizard of Leica AF software. Generally, the middle plane (or brightest z-plane) of a single BMC was selected (x25 zoom, 15.5×15.5 μm). A bleach area (region of interest ROI) of 20×20 pixels within the selected BMC was set for bleaching. We used the same ROI size for all our FRAP experiments. Photobleaching was done using a 405nm argon laser at 60% intensity for 6 frames. The bleaching session was preceded by 2 prebleach frames every 1.293s. Fluorescence recovery was images through 2 postbleach session: one with 10 frames every 3 seconds and a second one with 78 frames every 5 seconds. BMCs photobleaching has to reach at least 50% of fluorescence intensity to be used for analysis. For normalization purpose, two other ROIs were selected to account for overall fluorescence bleaching during the postbleach recovery, and to account for any background fluorescence. A non-fluorescent, background region (ROI2) and the area of total fluorescence (ROI3) were measured for each cell analyzed (Giakoumakis et al., 2017). Raw data from ROI1, ROI2 and ROI3 for each FRAP session were extracted and uploaded into EasyFRAP standalone version (versatile and public tool that assists quantitative and qualitative analysis of FRAP data, (Rapsomaniki et al., 2012)), double normalized, and then plotted using GraphPad Prism with standard error to the mean (SEM). FRAP experiments were considered successful and used for analysis when the gap ratio (total fluorescence remaining after bleaching) was higher than 0.7, and the bleaching depth (sufficient bleaching of the BMC in ROI1) was higher than 0.6 (meaning at least 60% bleaching from the initial intensity). The fluorescence recoveries were fitted using a double term exponential equation as previously reported to be most suitable for these FRAP recoveries (Rapsomaniki et al., 2012). The R square value of each fit had to be higher than 0.7 to be considered for analysis.

### Statistical analysis

The software GraphPad Prism 9 (9.2.0) was used for statistical analyses. Shapiro-Wilk normality test has been used to assess the plotted data for a normal distribution. Differences between the means were evaluated using ANOVAs and Holm-Sidak’s multiple comparison test for Gaussian distribution and t-test and Dunn’s multiple comparison test for non-Gaussian distribution. For all statistical tests, significance was taken as *\*p*< 0.05. Unless otherwise indicated, data values are presented as the mean ± standard error of the mean (SEM). Graphs were generated in GraphPad Prism 9 (9.2.0).

## ACKNOWLEDGEMENT

We wish to thank the Snow Foundation for their generous support towards establishing the transgenic zebrafish facility at Macquarie University and continued support of the researchers. We also wish to thank the zebrafish facility staff (past and present) for assistance in zebrafish care. This work was supported by the ALS Foundation Netherlands, FightMND, MNDRA, a Snow Foundation Fellowship, and donations towards the MND research at Macquarie University.

