## Supplementary Figures 1 to 5 for "The post-translational modification SUMO affects TDP-43 phase separation, compartmentalization, and aggregation in a zebrafish model"

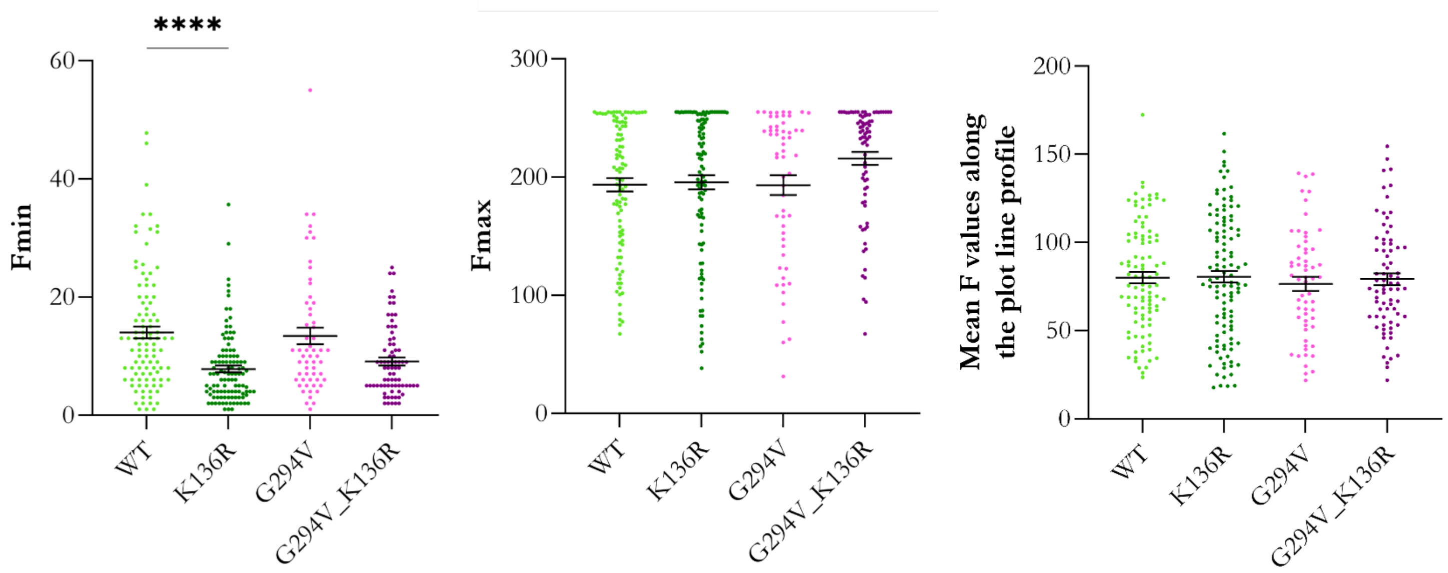


**Supplementary Figure 1**. Scatter plots illustrating fluorescence intensities (Fmin (cytoplasm), Fmax (nucleus), and average intensity values) from the plot line profile analysis. Fmin: 14.02 ± 1.00 for $TDP43^{WT}$, 7.85 ± 0.57 for $TDP43^{K136R}$, 13.42 ± 1.38 for $TDP43^{G294V}$ and 9.10 ± 0.67 for $TDP43^{G294V\_K136R}$, Kruskall-Wallis, n=57-109 neurons in 12-24 larvae, ****p<0.0001. Fmax: 193.6 ± 5.6 for $TDP43^{WT}$, 195.6 ± 5.9 for $TDP43^{K136R}$, 193.2 ± 8.3 for $TDP43^{G294V}$ and 215.8 ± 5.5 for $TDP43^{G294V\_K136R}$, Kruskall-Wallis, n=57-109 neurons in 12-24 larvae, ns. Average intensity values measured among the plot line profile analysis: 80.1 ± 3.1 for $TDP43^{WT}$, 80.6 ± 3.4 for $TDP43^{K136R}$, 76.6 ± 4 for $TDP43^{G294V}$ and 79.2 ± 3.4 for $TDP43^{G294V\_K136R}$, Kruskall-Wallis, n=57-109 neurons in 12-24 larvae, ns.

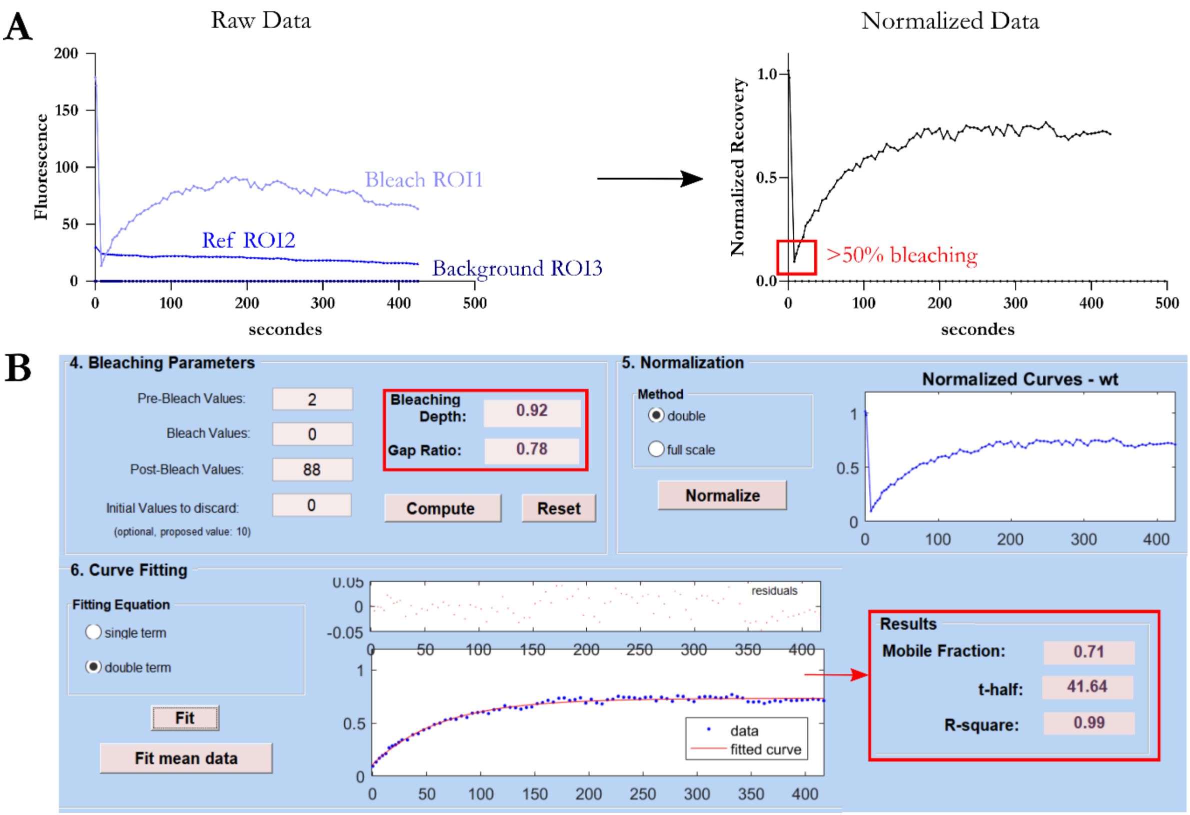


**Supplementary Figure 2. (A)** Raw data of fluorescence intensity collected from the 3 regions of interest (bleach, reference, and background) during confocal imaging session post and pre-bleach. As explained in the methods, ROI2 and ROI3 were used to normalized ROI1 data and plotted to observe the recovery after photobleaching. Bleaching over 50% was one of the criteria for further analysis with the EasyFRAP software. **(B)** EasyFRAP interface illustrating the workflow and curve fitting parameters. Criteria for inclusion in calculating the mean fluorescence recovery included a bleaching depth >0.6 and a gap ratio >0.7 (4. Bleaching parameters). After double normalization (5. Normalization), data were curve fitted using double term equation (6. Curve Fitting). The mobile fraction, t-half time, and R-square values were extracted, plotted, and analyzed using GraphPad Prism.

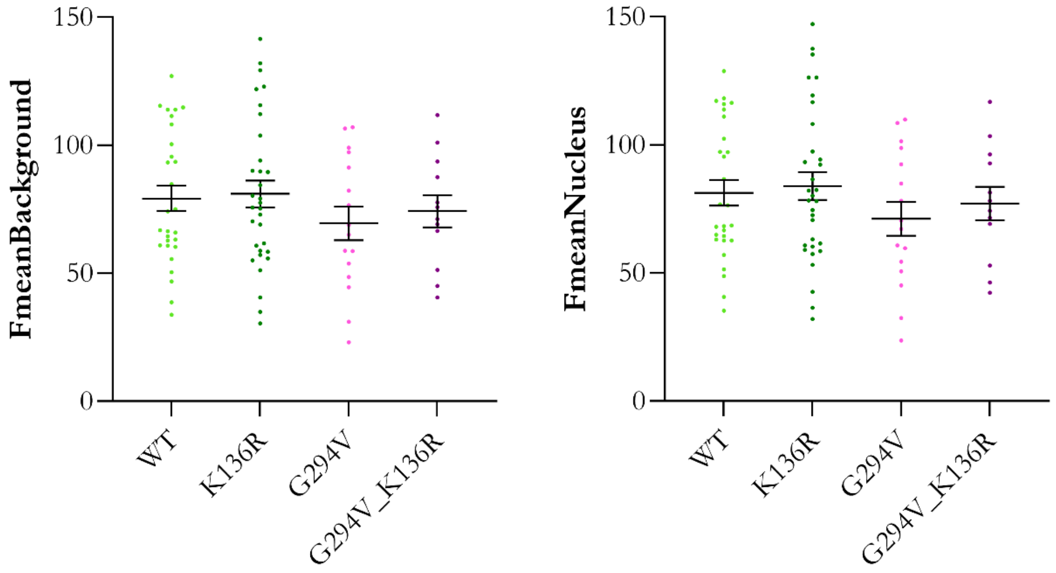


**Supplementary Figure 3.** Scatter plots illustrating the analysis of diffuse TDP-43 in the background of the nucleus. FmeanBackground and FmeanNucleus data are all non-significantly different. FmeanBackground of 79.2 ± 4.9 for WT, 81 ± 5.2 for K136R, 69.5 ± 6.6 for G294V, 74.3 ± 6.3 for G294V_K136R, One-way ANOVA, n=12-32, ns, and FmeanNucleus of 81.3 ± 5 for WT, 83.9 ± 5.4 for K136R, 71.2 ± 6.7 for G294V, 77.1 ± 6.6 for G294V_K136R, One-way ANOVA, n=12-32, ns.

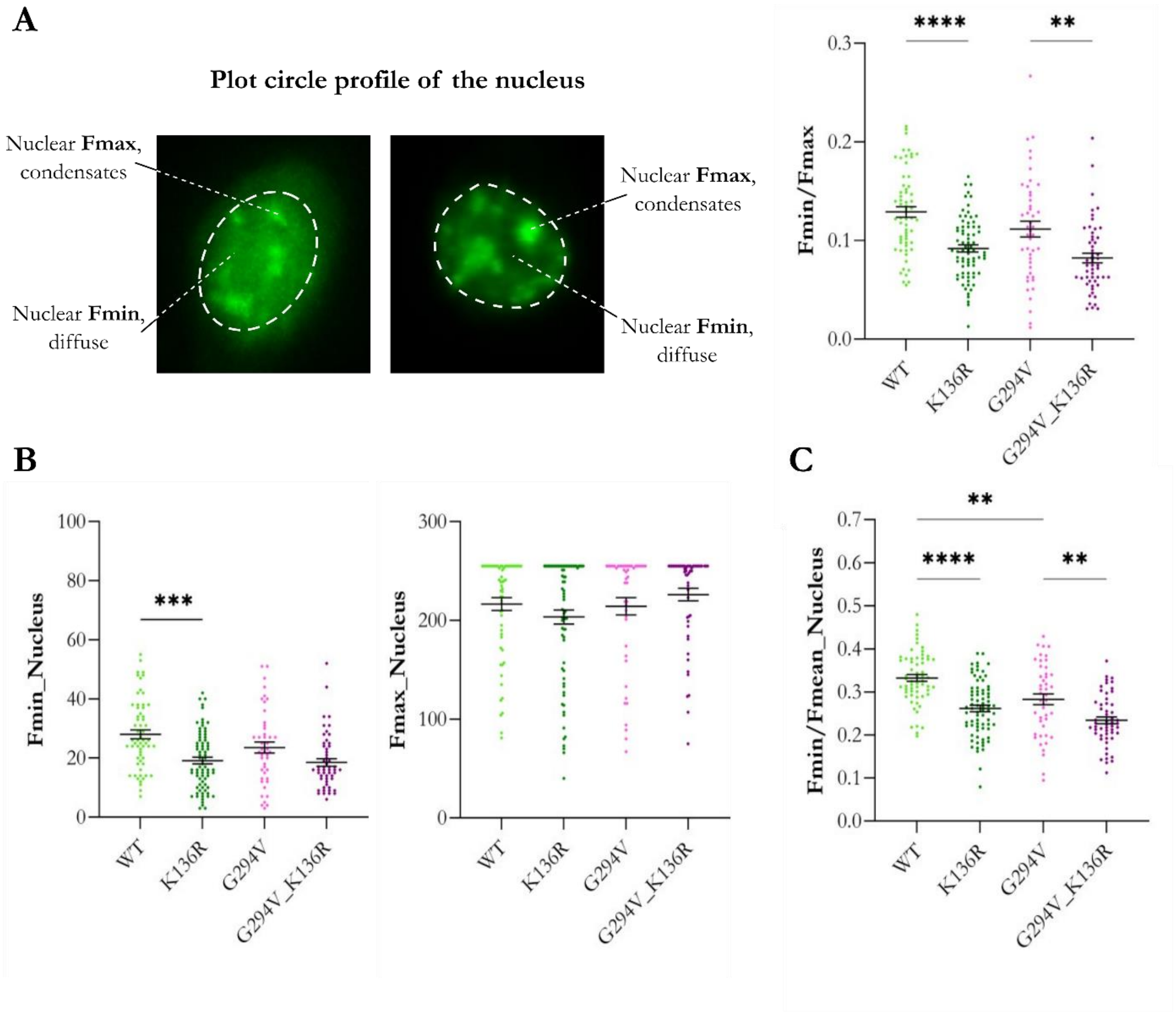


**Supplementary Figure 4**. **(A)** Representative images showing the plot circle profile of the nucleus. Fmin values represent non-condensated TDP-43 while Fmax represents bright condensates. Fmin/Fmax of 0.129 ± 0.005 for TDP43$^{WT}$, 0.092 ± 0.004 for TDP43$^{K136R}$, 0.112 ± 0.008 for TDP43$^{G294V}$ and 0.082 ± 0.005 for TDP43$^{G294V_K136R}$, Kruskal-Wallis, n=45-75, ****p<0.0001 **p=0.037. **(B)** Quantitative comparison of Fmin and Fmax values. Fmin of 28.1 ± 1.5 for TDP43$^{WT}$, 19.1 ± 1.1 for TDP43$^{K136R}$, 23.6 ± 1.9 for TDP43$^{G294V}$, and 18.5 ± 1.3 for TDP43$^{G294V_K136R}$, Kruskal-Wallis, n=45-75, ***p=0.0001. Fmax didn't show any differences. **(C)** Ratios of diffuse TDP-43 (Fmin) and the overall average fluorescence intensities in the nucleus. Fmin/FmeanNucleus of 0.332 ± 0.007 for TDP43$^{WT}$, 0.262 ± 0.007 for TDP43$^{K136R}$, 0.283 ± 0.013 for TDP43$^{G294V}$ and 0.233 ± 0.008 for TDP43$^{G294V_K136R}$, ANOVA, n=45-75, ****p<0.0001 **0.0012 < p < 0.0020. All FmeanNucleus data are non-significantly different (data not shown).

MSEYIRVTEDENDEPIEIPSEDDGTVLLSTVTAQFPGACGLRYRNPVSQCMRGVRLVEGILH

APDAGWGNLVYVVNYPKDNKRKMDETDASSAVKVKRAVQKTSDLIVLGLPWKTTEQDLKEYF RMM1

STFGEVLMVQVKKDLKTGHSKGFGFVRFTEYETQVKVMSQRHMIDGRWCDCKLPNSKQSQDE

PLRSRKVFVGRCTEDMTEDELREFFSQYGDVMDVFIPKPFRAFAFVTFADDQIAQSLCGEDL RMM2

IIKGISVHISNAEPKHNSNRQLERSGRFGGNPGGFGNQGGFGNSRGGGAGLGNNQGSNMGGG LCD

MNFGAFSINPAMMAAAQAALQSSWGMMGMLASQQNQSGPSGNNQNQGNMQREPNQAFGSGNN

SYSGSNSGAAIGWGSASNAGSGSGFNGGFGSSMDSKSSGWGM

**Supplementary Figure 5**. **N-terminal and C-terminal part forms a network to tune phase separation considering TDP-43 amino acid sequence.** Arginine and tyrosine (bold R and Y) positively charged residues are modulators of phase separation. RGG motif in orange is on crucial signature motif for LLPS. All underlined residues are post-translationally modified according to PhosphoSite. Important domains for LLPS are represented in turquoise for the first RNA Recognition Motif RRM1, blue for the second RRM2 and in red for the low complexity domain LCD in the C-terminal part of TDP-43.

**Supplementary Video 1.** Videos representing dynamic condensates moving inside the nucleus.

**Supplementary Video 2.** Videos representing fission and fusion events of condensates inside the nucleus.
